# Individual differences in theta-band oscillations in a spatial memory network revealed by EEG predict rapid place learning

**DOI:** 10.1101/2020.06.05.134346

**Authors:** Markus Bauer, Matthew G. Buckley, Tobias Bast

## Abstract

Spatial memory has been closely related to the medial temporal lobe (MTL), and theta-oscillations are thought to play a key role. However, it remains difficult to investigate medio-temporal lobe (MTL) activation related to spatial memory with non-invasive electrophysiological methods in humans.

Here, we combined the virtual delayed-matching-to-place (DMP) task, reverse-translated from the watermaze DMP task in rats, with high-density electroencephalography (EEG) recordings. Healthy young volunteers performed this computerised task in a virtual circular arena, which contained a hidden target whose location moved to a new place every four trials, allowing the assessment of rapid memory formation.

Using behavioural measures as predictor variables for source reconstructed frequency specific EEG power, we found that inter-individual differences in ‘search preference’ during ‘probe trials’, a measure of 1-trial place learning known from rodent studies to be particularly hippocampus dependent, correlated predominantly with distinct theta-band oscillations (approx. 7 Hz), particularly in the right temporal lobe, the right striatum and inferior occipital cortex or cerebellum. Notably, this pattern was found with very high consistency during both encoding and retrieval/expression, but not in control analyses and could not be explained by motor confounds. Alpha-activity in sensorimotor and parietal cortex contralateral to the hand used for navigation also correlated with search preference, which likely reflected movement-related factors associated with task performance.

Relating inter-individual differences in ongoing brain activity to behaviour in a continuous rapid place learning task that is suitable for a variety of populations, we could demonstrate that memory related theta-band activity in temporal lobe can be measured with EEG recordings, revealing a presumed network of MTL, striatum and cerebellum and/or inferior occipital cortex that may interact through theta oscillations. This approach holds great potential for further studies investigating the interactions within this network during encoding and retrieval, as well as neuromodulatory impacts and age-related changes.

## Introduction

A large body of research in humans and animal models supports that the hippocampus and the surrounding medial temporal lobe (MTL) cortices play a crucial role in certain types of memory, including place memory (Burgess et al., 2002; Bast, 2007; Morris, 2007). Theta oscillations are a prominent neural activity pattern recorded from MTL regions, in particular the hippocampus, especially during movements in real and virtual environments. Theta oscillations have been associated with encoding and retrieval of memory, including place memory, and this has been suggested to reflect that theta oscillations facilitate underlying synaptic plasticity mechanisms, separate encoding and retrieval processes, or coordinate the activity of neuronal ensembles that are distributed across hippocampus and other brain regions and interact to support memory encoding, retrieval or expression (O’Keefe and Burgess, 1999; Kahana et al., 1999; Cornwell et al., 2008; Colgin, 2013; Hasselmo and Stern, 2014; Herweg et al., 2020).

Watermaze tests of place learning and memory in rodents, and corresponding reverse-translated human paradigms in real or virtual environments, have long been used to study hippocampal function (Morris et al., 1982; Morris, 2007; Cornwell et al., 2008; Pu et al., 2017; Buckley and Bast, 2018). In common variants, the rodent or human participant has to find a hidden goal that remains in the same place over many trials, allowing for incremental learning of the place with reference to distal cues surrounding a circular, featureless maze. Place memory of where the goal is located in relation to the distal cues (i.e., allocentric place memory) is reflected by relatively short latencies and direct paths to the goal when the animals are placed into the maze from different start positions (which discourages use of egocentric strategies), and by search preference, i.e. persistent searching around the goal location, when the goal has been removed for a probe trial. Although incremental place learning performance depends on the hippocampus, rodent studies have shown that one-trial place learning performance, as measured using the delayed-matching-to-place (DMP) watermaze variant, is a more sensitive index of hippocampal function (reviewed in (Buckley and Bast, 2018)). The DMP task requires the continuous rapid updating of place memory, resembling the everyday task of remembering where we parked our car on a particular occasion. In a common DMP protocol (Steele and Morris, 1999; Bast et al., 2009), which we have recently reverse-translated into a virtual DMP task for human participants (Buckley and Bast, 2018), the goal moves to a new place every four trials. One-trial place learning on the rodent and human DMP task is reflected by marked reductions in latency and path lengths to the goal from trial 1 to 2, with little further improvements on subsequent trials, and by a marked search preference for the vicinity of the goal location when trial 2 is run as probe (when the goal is removed). Rodent studies have shown that such one-trial place learning performance is highly hippocampus-dependent, being more sensitive to disruption of hippocampal function than incremental place learning performance; for example, partial hippocampal lesions and manipulations of hippocampal synaptic plasticity, which can leave incremental place learning performance relatively intact, markedly impair one-trial place learning performance (reviewed in (Buckley and Bast, 2018)). Moreover, the DMP task allows for the repeated study of encoding and retrieval/expression of new place memory within the same participants. Thus, the virtual DMP task may be particularly suitable to reveal the neural oscillations associated with encoding or retrieval/expression of hippocampus-dependent place memory performance in human participants.

There is considerable inter-individual variance in cognitive performance parameters. For instance, ageing affects spatial memory in particular and this may happen through decline of cholinergic neuromodulation (Richter et al., 2014). In Alzheimer’s disease (AD), the MTL and the nucleus basalis are amongst the first brain areas to be damaged (Braak and Braak, 1995), and several studies have shown that a deficit in spatial memory in AD correlates with hippocampal damage (Miller, 1973; Kramer et al., 2004). We have previously shown that functional connectivity of the nucleus basalis to cortex can predict the therapeutic response to cholinesterase inhibitors by patients suffering from mild cognitive impairment; more specifically, functional connectivity of the nucleus basalis predicted the improvement of cognitive performance scores in response to treatment with cholinesterase-inhibitors (Meng et al., 2018). A question that arises is, how do such cholinergic changes mediate memory processes? One candidate mechanism is through hippocampal theta oscillations as these are known to be dependent on cholinergic input (Tiesinga et al., 2001; Newman et al., 2012). The vDMP task might therefore serve as an ideal model to study such mechanisms in the future.

In the present study, we aimed to examine whether electroencephalography (EEG) was suitable to measure memory related theta oscillations from the MTL in human participants performing the virtual DMP test. Previous studies have used magnetencephalography (MEG) to localize spatial navigation-related theta-oscillations (Cornwell et al., 2008; Kaplan et al., 2012; Kaplan et al., 2014; Pu et al., 2017), but MEG is far more expensive than EEG and these authors also used individual anatomical MRIs of the head to construct appropriate volume-conductor (and thus biophysical forward) models for source analysis, making this method much less commonly available to use for future studies of individual differences in the context of ageing or other clinically relevant conditions or in particular diagnostic tests for early dementia. Additionally, we aimed to perform a largely data-driven analysis to reveal key frequency bands (within the range of 2-15 Hz) and brain regions most robustly associated with performance, rather than constraining our analysis on predetermined frequencies or regions of interest.

## Materials and Methods

### Participants

Initially 26 participants were recruited, predominantly students from the University of Nottingham recruited through an opportunity sample as part of MSc summer research projects, in line with ethical guidelines and having given informed consent. Participants were given an inconvenience allowance for their participation. The sample size was chosen to represent a compromise between statistical power and practical considerations. It was a little higher than sample sizes in previous MEG studies of spatial memory and navigation related theta (18 participants (Pu et al., 2017); 19 participants (Kaplan et al., 2012)). The data of five participants had to be excluded due to problems with their EEG data, either trigger problems or electrical noise issues caused by an electrical device on the roof of the semi-shielded room in which the experiment was conducted. The data of 21 participants were thus included in this report (mean age: 24.4 years, std-dev: 2.9 years).

### Virtual DMP task

The virtual DMP task was run using MazeSuite Software (www.mazesuite.com) (Ayaz et al., 2008) on a Windows 7 computer, using procedures adapted from our previous study (Buckley and Bast, 2018). Given that we have described the environment used in the current study in detail elsewhere (Buckley and Bast, 2018), we give only a brief overview of the materials to aid understanding of the task, and focus instead on the procedural details for the present EEG study.

The virtual environment was identical to that used in our previous study, and comprised of a circular grass lawn surrounded by a brown fence, beyond which eight cues (a tree, hot air balloon, star, plane, tower, windmill, satellite and a planet), could be seen by participants and could be used for spatial orientation. These cues were varying distances from the fence, but distributed at regular 45 degree intervals around the circular fence (**Figure 1A,B**).

**Figure 1:**
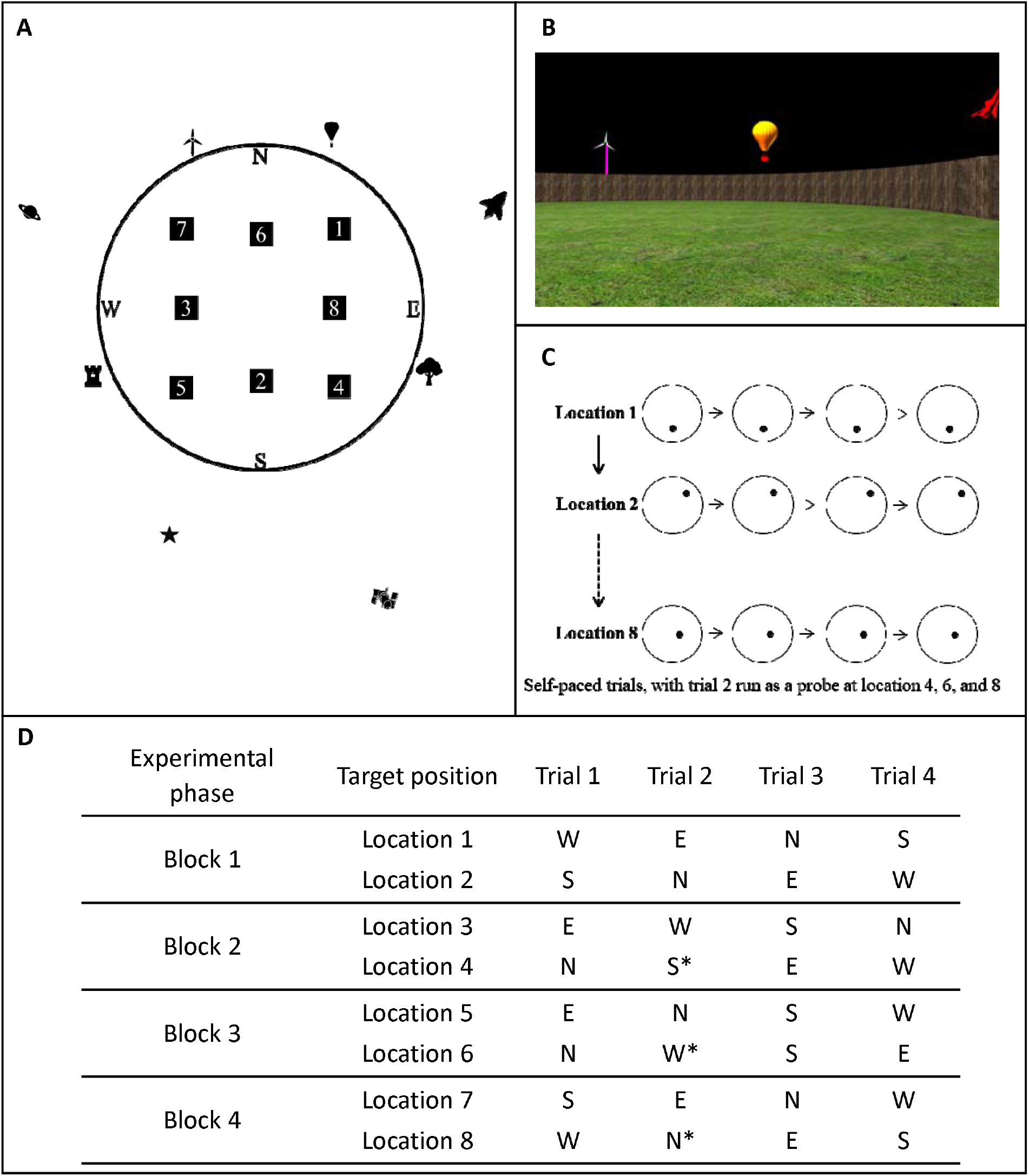
The virtual DMP task and the structure of the experimental sessions. **A)** and **B)** Participants were placed within a circular environment, with landmarks presented at varying distances from the circular wall, and were instructed to find a hidden target (see Fig. 2 for an illustration of search paths). **C)** In order to repeatedly test rapid place learning, the hidden goal (filled circles in panel C) moved after every 4 trials, and was placed at 8 different locations during an experimental session (filled squares in panel A). **D)** Each participant completed one experimental session split into 4 blocks. Each block started with a 2 min resting state period, followed by two sets of 4 navigation trials. During each set of 4 trials, the goal was placed in the same location, and participants began one trial from each of the notional cardinal points of the environment. Participants were then instructed that the goal had moved, and completed another set of 4 trials navigating to the new location. In trial 1, participants could not know the location of the target and had to search for it. In trials 2-4, the participants could use the memory of the location acquired during trial 1 in order to navigate to the target efficiently. At locations 4, 6, and 8, the second of the four navigation trials was ran as a probe trial (marked by an *), during which no feedback was given when participants crossed the target location. Probe trials continued for 60 s, during which participants’ “search preference” for correct location could be measured. EEG was recorded continuously throughout the entire session.

The virtual arena was viewed from a first-person perspective, and participants were instructed to control movement through the environment using the four cursor keys on the keyboard with their right hand. Presses on the ‘up’, ‘down’, ‘left’, and ‘right’ keys permitted the participant to move forwards, backwards, and to turn (rotate) counter-clockwise and clockwise, respectively. Trials started with the participant being placed in the environment facing the perimeter fence. As described elsewhere (Buckley and Bast, 2018), the environment was modelled on a rodent watermaze (Bast et al., 2009) and, thus, travelling from the start location to the opposite side of the fence (i.e. the diameter of the circle) took approximately 20 s.

During an experimental session, participants were instructed to search for an invisible target, ‘William the Worm’, a cover story that has been used successfully with children and adults of varying ages (Buckley et al., 2015; Buckley and Bast, 2018), and to remember the location of the target using the cues arranged around the fence. Each participant completed one session of 32 trials, in which William the Worm would be located at eight different locations, for four trials at each location (**Figure 1C**). To prevent the use of egocentric strategies to navigate to each target location, participants began each of the four trials to the same target location at one of four different start positions spaced evenly along the fence perimeter (the notional N, E, S, and W of the environment). The sequence of start positions and target locations was the same for all participants.

The overall 32 trials (to 8 different locations) that each participant completed were split equally over four experimental blocks (**Figure 1D**). Each block started with a 120 s resting state period to serve as a baseline for the EEG analysis, in which a fixation cross was shown on a black screen throughout. Subsequently, within each block, participants would find William the Worm at two different locations. For each location within a session, the participant was given four consecutive trials to find the target (in the same location), before they were notified that the location of the target had changed

For most trials of the experiment participants were either given feedback once they found the target, or the target was highlighted to them after 120 s by a white flag appearing at the goal location. Having navigated to the target location, participants received a congratulatory message (‘Congratulations!!! You found William! Congratulations!!!’) and initiated the next trial using the Enter key. However, in experimental blocks 2-4, probe trials were administered on the second trial at the second location. Here, unlike every other trial in the task, participants were not informed (and the trial did not end) when they crossed the target location. Instead, participants were allowed to move freely for 60 s in order to allow the measurement of ‘search preference’ (see in Behavioural data analysis). Each probe trial ended after 60 s had elapsed, after which a message (‘Keep looking for William!’) was displayed on the screen for 1 s, and then the next normal trial (i.e., Trial 3) began automatically.

### Behavioural data analysis

All data analysis was carried out in MATLAB. The time-series of participants’ positions were therefore imported and temporally aligned with the recorded EEG signals. For primary analysis we calculated the following path-length measures: 1) ‘total path length’ of their translational (not rotational) movements in planar Euclidean space during the trial whilst searching for the target, 2) ‘path efficiency’ as the ratio between the ‘total path length’ and the distance of their starting position to the target position, which varies substantially from trial to trial (see Figure 2). Data from the first location (i.e. the first four trials) were excluded from analysis since participants were still adapting to the task.

**Figure 2:**
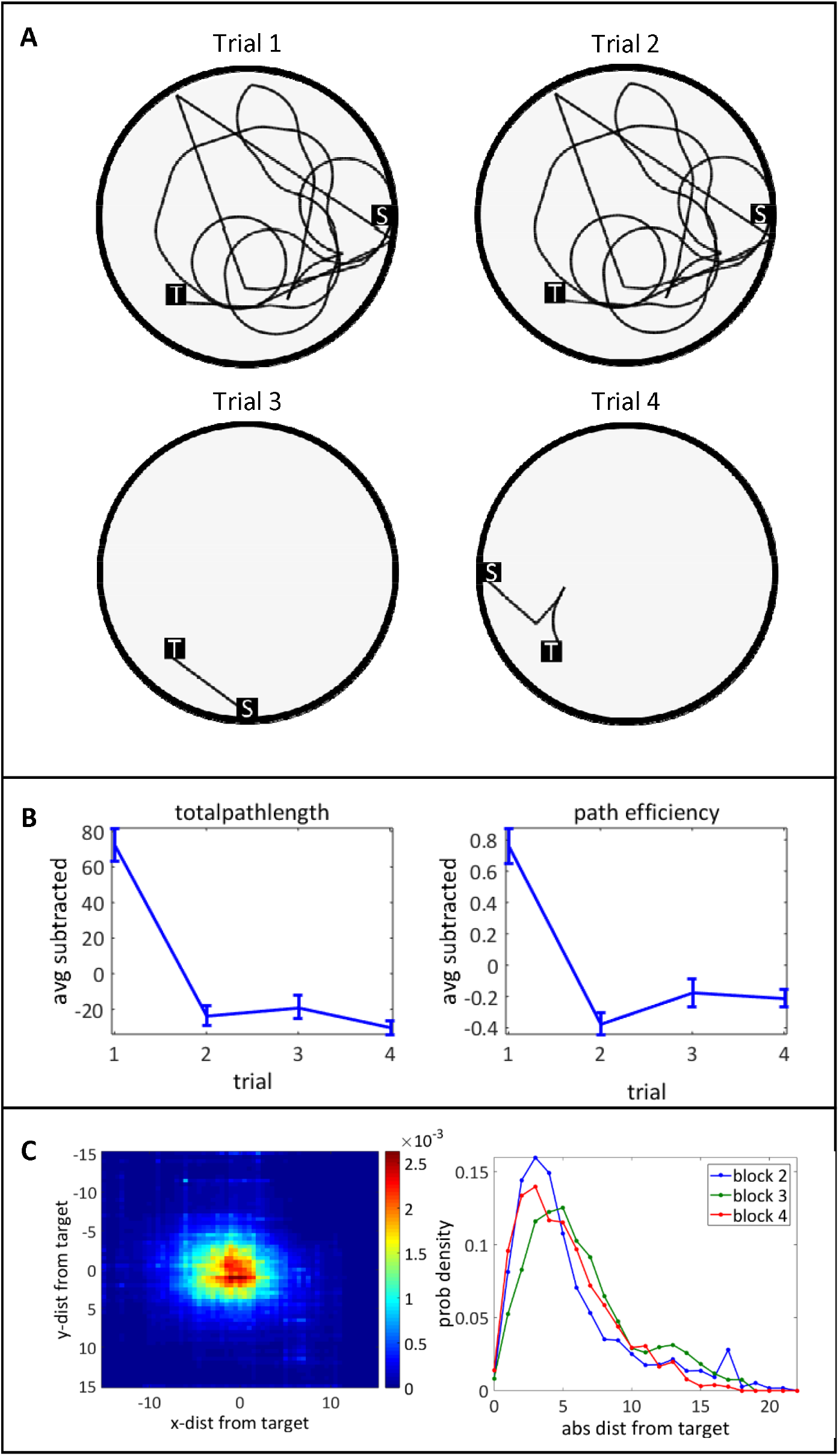
Behavioural performance on the virtual DMP test. **A)** Illustration of one participant’s search paths during the four trials to one location. **B)** Path-length measures as a function of trial, averaged across all locations and participants. Left: Total path length is the entire translational distance (without rotations) travelled by the participant; numbers represent (arbitrary) ‘maze units’ as distance measure. Right: Path efficiency is path length normalized to the distance of starting position to target location. Averages of each participant across trials have been subtracted; error-bars thus represent standard error of mean condition differences across participants, i.e. confidence intervals. **C)** Left: Average heat map of participant locations during probe trials, with the target location at 0,0 (probability density across participants and all probe trials). Right: Histogram of locations (average probability density) for the three different probe trials in sequence of occurrence. Numbers represent ‘maze-units’ as spatial dimension.

In addition, we calculated a ‘search preference’ measure, which has been shown to be particularly dependent on hippocampal function in rat studies using the watermaze DMP test (Bast et al., 2009; Pezze and Bast, 2012; McGarrity et al., 2017). In this study, we measured the ‘search preference’ for the correct location during the probe trial as the average distance the participant kept from the target throughout the probe trial (with lower average distance reflecting higher search preference). A similar measure has been used in previous watermaze studies in rodents (Gallagher et al., 2015), although a more common measure of search preference in rodent watermaze and human virtual maze studies is the time the subjects spent within a pre-defined region around the target, often expressed as a proportion of time spent in other parts of the maze (Buckley and Bast, 2018). The reasons we used the distance-to-target measure (which was highly correlated with the proportion of time spent near the target) in the present study were 1) that it only depends on the target location, without the requirement to define a zone around it or comparison areas, and 2) ease of implementation.

### EEG analysis

All analysis was conducted using FieldTrip (Oostenveld et al., 2011) and custom-written analysis software in MATLAB. Continuous data from both the rest and maze periods was notch-filtered for power line noise at 50 Hz, temporally aligned to the maze-data and then split into 1-s epochs (pseudo-trials) to facilitate the analysis. All data from one block (of which there were four per participant) were concatenated and checked for excessive artefacts using a summary statistic (total power calculated for each trial and electrode) provided by ft_rejectvisual. A principal component analysis (PCA) was then computed on these data and eye-blink components and obvious artefact components were removed from the data. A final inspection of any residual artefacts was carried out, using, again, the summary statistic provided by ft_rejectvisual.

In order to have a reasonable separation of signals from different sources, e.g. motor cortex, temporal lobe and occipital cortex, we conducted all data analysis in the source space using a beamformer transformation (Van Veen et al., 1997; Gross et al., 2001). To this end, we used the boundary element volume-conductor model (BEM) of the segmented standard MNI brain as implemented in the Fieldtrip toolbox (Oostenveld et al., 2011; Oostenveld et al., 2003). Average digitised electrode positions were aligned to the head surface (based on MNI brain) and the leadfields were calculated for a three-dimensional grid covering the entire (MNI) brain with a spatial resolution of 1cm.

To calculate the source reconstructed power spectra with a good signal to noise ratio and in a computationally efficient way, we used the following approach: A time-domain linearly constrained maximum variance beamformer (Van Veen et al., 1997) was calculated on low-pass filtered (cutoff 30 Hz) data from all epochs including the resting state period, using the covariance matrix from all these epochs concatenated across the four blocks and the leadfield matrix. This approach gives particularly robust filters (Litvak et al., 2010; Bauer et al., 2012b; Bauer et al., 2014). To obtain source estimates of power spectral density estimates, we then projected the real part (not the imaginary) of the (‘sensor-level’) cross-spectral-density matrix (calculated in 1Hz steps over 1s epochs using a Hanning taper), averaged for all epochs of a particular trial type (resting state, Trial 1, Trial 2 exclusive probe trials, Trial 3, Trial 4 and probe trial) through the beamformer.

This gave thus trial specific estimates of power-spectral-density (2-15Hz in steps of 1Hz) for each of 2015 grid points in the brain (1cm regular grid) – separately for resting state, Trial 1, Trial 2, Trial 3, Trial 4 and probe trials, averaged across locations. Regarding the choice of spatial resolution and frequency range, see the next paragraph.

### Statistical analyses of EEG correlates of behaviour

The mere comparison of the navigation trials on the virtual DMP task vs rest is confounded by motor activations due to button presses, as well as dynamic visual input and enhanced attentional load during the navigation trials. Hence, this comparison is of limited suitability to reveal any memory related activations.

In order to do so, we sought to correlate EEG power spectral density estimates with behavioural performance measures in the task. We chose the following approaches: 1) Use the individual difference in path efficiency between trial 2 and trial 1 as a predictor of individual EEG activity during trial 1 to potentially reveal encoding processes; 2) use the same behavioural measure as a predictor of individual EEG activity during trial 2 to reveal retrieval/expression processes; 3) use the search preference for the target location, as reflected by average distance to target, during probe trials as predictor of EEG activity in probe trials to reveal retrieval/expression processes; and 4) use the same behavioural variable (average distance to target during probe trial) to predict EEG activity during trial 1 to reveal encoding processes. Further control analyses following the same logic are described in the results section.

We aimed to perform a data-driven analysis rather than aiming to confirm a priori hypotheses and thus constraining our analysis on particular frequencies or regions of interest (also given the leakage of spatial filters, particularly with a canonical forward model). Given the large number of grid points (or ‘voxels’) covering the brain and the multiple frequencies involved, we corrected for the multiple comparison problem (for mass-univariate testing), using a cluster permutation approach (Maris and Oostenveld, 2007).

The frequency range and spatial resolution were chosen with the following rationale:

1. to provide a compromise between statistical sensitivity (power) when correcting for multiple comparisons with mass-univariate analyses and to cover a reasonable range with sufficient specificity;
2. regarding the beamformer spatial resolution, although 1cm is on the lower side, the use of a canonical volume conductor model (in contrast to individual ones based on segmented individual MRIs of participants’ brains) and regularization of the covariance matrix would likely prevent a higher effective spatial resolution anyway;
3. regarding the frequency range, the lower bound was chosen to be just above the Rayleigh frequency, whereas the upper bound was chosen as to encompass a reasonably broad range of frequencies, including and extending beyond the low-frequency MTL oscillations that have been implicated in spatial navigation and memory (Ekstrom and Watrous, 2014) and that were the focus of the present study, without making the statistical tests too insensitive.

Mass-univariate regression analyses were conducted using the behavioural metrics as a predictor variable and the EEG power of the particular trial type as the criterion with individual participants and blocks being units of observation; within and between-subject variance was thus pooled in this approach to harvest as much of behavioural variation as possible. Since relatively large amplitude differences existed between participants, possibly related to impedances, the data were a priori normalized using the following approach: the individual (participant specific) source estimates of power spectral density were divided by the (individual) squared average of the square root of all power spectral density estimates across all frequencies and grid points during the 2 min resting state periods prior to each block.

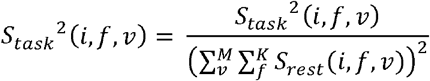

where *i* is the index to the participant, *f* the index for *K* frequencies and *v* the index for *M* grid-points (or voxels) and S^2^ is the power spectral density estimate (i.e. the squared estimate of the complex Fourier coefficient). The cluster-permutation approach (Maris and Oostenveld, 2007), briefly summarised, was applied as follows: spatio-spectral clusters (in three spatial plus a spectral dimension) were formed of adjacent data-points that survived the alpha-threshold of p<0.01 for two-sided testing on the regression coefficient. The cumulative t-value of the regression coefficients within each cluster was then calculated. This calculation for the real data was then repeated for 1000 randomised data-sets, where, for each combination of frequency bin and spatial grid point (= ‘voxel’), the associated behavioural parameter was shuffled randomly across observations (participants). A histogram of the maximum cluster metric for each randomization was formed, serving as the reference, or Null-distribution. Those clusters of the real data for which the aggregated statistic was above the 5% percentile of this reference distribution, the omnibus null-hypothesis can be rejected.

## Results

### Behavioural data: one-trial place learning on the virtual DMP test

**Figure 2A** shows examples of the search paths taken by one participant during the four trials to one location. On trial 1, the participant searches for the new (unknown) target location, in this example in a mainly circular pattern (for other search strategies during trial 1, see Buckley and Bast, 2018, Fig. 2). In subsequent trials, this search path is typically markedly more direct (i.e., shorter), reflecting 1-trial place learning and leading to the behavioural summary statistics shown in **Figure 2B, left.** What can also be appreciated from **Figure 2A** is that the distance of initial starting position in the maze to target varies markedly across trials, target locations and blocks (although this is kept constant across participants). To remove this variability, total path length was normalised to this initial distance and presented as ‘path efficiency’ (**Figure 2B, right**). The latter metric effectively expresses the ‘straightness’ of the path taken to the target. Total path length was significantly shortened with higher trial numbers (F(3,20) = 47.85, p < 10^-8^), as was path-efficiency (F(3,20) = 24.45, p <10^-6^), with a sharp reduction from trial 1 to 2, reflecting 1-trial place learning. For all measures, the average of each participant across conditions was first subtracted to make the SEMs interpretable regarding statistically significant differences (within-subject design).

When the second trial to a new location was run as a probe trial, where participants roamed around for 60 s without being given feedback about the target location, participants typically showed marked search preference for the target location by moving around in close proximity to the target (**Figure 1C**), reflecting 1-trial place learning. Overall, both path length measures and the search preference measure revealed marked 1-trial place learning, similar to previous DMP studies in rodents and humans (Buckley and Bast, 2018).

### Correlations between behaviour and EEG activity

Since the comparison between maze trials 1-4 on the one hand and rest periods on the other hand confounds mnemonic processes with motor outputs, visual input and attentional demands, we focused on the analysis of inter-individual differences in behaviour and EEG parameters. All analyses were performed in three-dimensional source space, using a regular 1cm grid, based on a canonical forward model (MNI brain).

### Reductions in path-length measures from T1 to T2 show no significant correlations with EEG activity

As a first step, we used the difference in path length and in path efficiency, respectively, between trial 1 and trial 2 as a predictor variable for EEG activity in either trials 1 (encoding) or in trials 2 (retrieval/expression, where these were not probe trials). The cluster analysis, correcting for multiple comparison across space and frequency yielded no results that crossed the significance threshold (all p > 0.3), both for trials 1 and trials 2. Moreover, lowering the univariate clustering-threshold to p<0.05, did not change this result. We interpret this to be due to the comparably small variability in the spatial distance measures (see **Figure 2B**).

### Negative correlations of theta oscillatory activity with distance to target – potential correlates of encoding and retrieval/expression processes

We next turned to the ‘average distance to the target’ during the ‘probe trial’, reflecting ‘search preference’ for the target location, which has been shown to be particularly dependent on hippocampal function in rat studies (Bast et al., 2009; Pezze and Bast, 2012; McGarrity et al., 2017). We used the average distance to the target during the three probe trials as a predictor variable for EEG data during both the same probe trial itself (i.e., reflecting retrieval/expression processes), as well as for the EEG data during the corresponding trial 1 (i.e., reflecting the encoding process). This analysis revealed negative correlations of oscillatory activity during probe trials (retrieval/expression) and during trial 1 (encoding) with average distance to target during probe trials (i.e., oscillatory activity predicted higher search preference), which were specific with respect to frequency and brain region (**Figure 3 A-D**).

**Figure 3:**
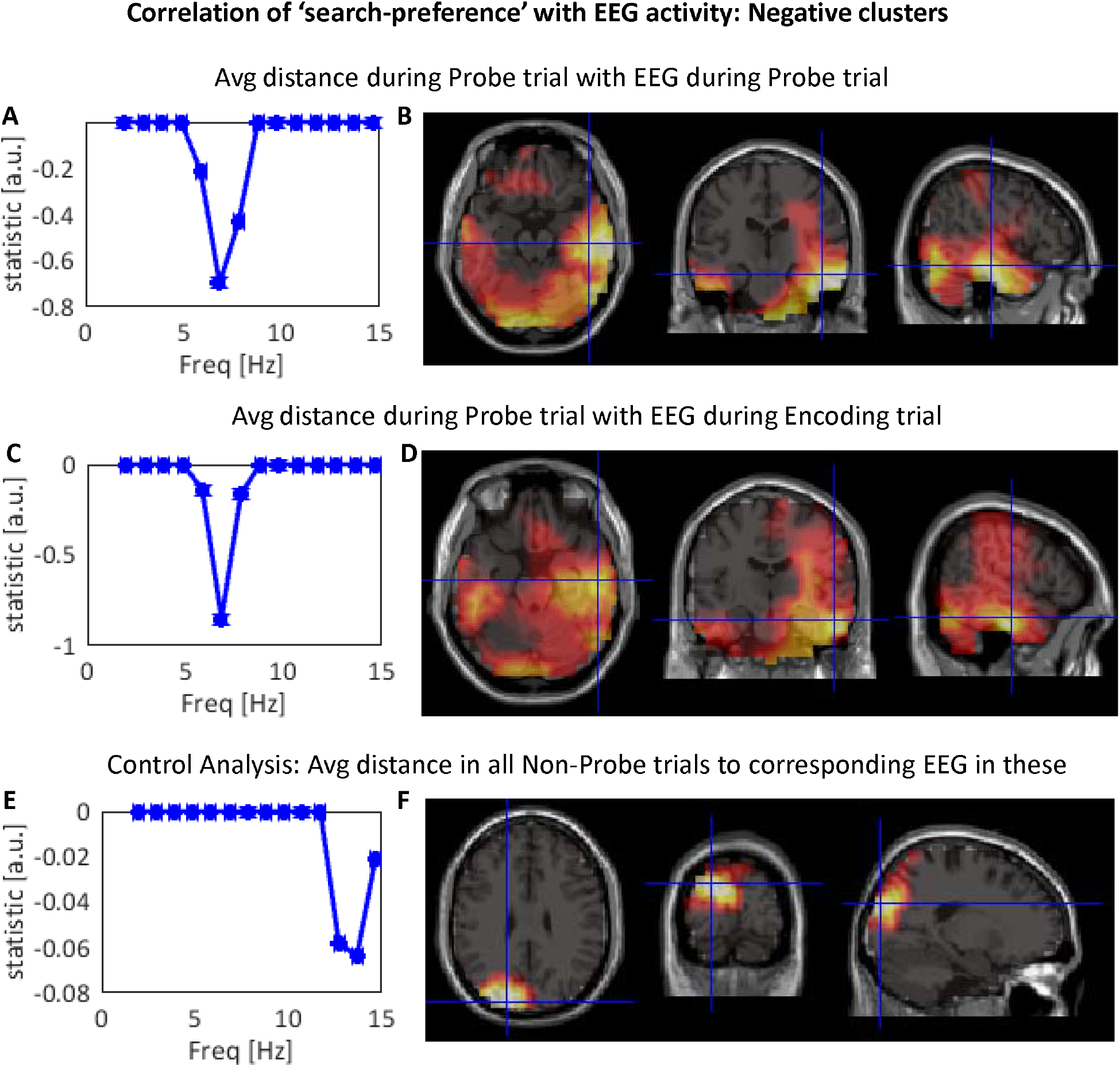
Negative correlations between oscillatory EEG activity and distance to target. **A)-D)** Negative correlations of EEG activity during probe trials (retrieval/expression) and trial 1 (encoding) with average distance to target during probe trial. Crosshairs in brain topographies show the location of cross-section planes (chosen to provide good overview). **A)** Frequency spectrum of the omnibus significant cluster for the probe trial, correcting for multiple comparisons across frequencies and 3 spatial dimensions; higher theta-band activity during the probe trial is correlated with smaller distance to target. **B)** Axial, coronal and sagittal views of the thresholded cluster (brain activations are sign-inverted to frequency spectra): activation is centred in (predominantly right) temporal cortex, inferior occipital cortex or cerebellum and striatum, forming one large cluster. **C)-D)** Same as A)-B), but now for the oscillatory activity during trial 1 (encoding); note the spectral and spatial similarity of the effects. **E)-F)** As a control analysis, the average distance during all non-probe trials was correlated to the EEG activity in the corresponding trials (to test for potential confounds, e.g. related to sensori-motor factors). The largest cluster of this analysis was a peak in occipital alpha-/beta activity (also sign-inverted) that did, however, not reach omnibus significance.

**Figure 3A** shows the spectrum of the thresholded statistics for the correlation between probe trial EEG activity and average distance to target during the probe trials, i.e. during retrieval/expression. Only data points that ‘survived’ the omnibus significance threshold of p<0.05 (two-sided) on the cluster level have values above zero, weighted by the number of data bins that reached significance – i.e., the strength of the spectral statistics depends on the effect size in the respective frequency bin, as well as on the number of spatial grid points that are significant at that particular frequency, and thus reflects a weighted average. There was a negative theta peak (6-8 Hz), indicating that enhanced theta power correlates with a smaller average distance to the target, and thus more accurate 1-trial place memory performance. **Figure 3B** shows the spatial signature of this cluster (significant at p < 0.05, two-sided and corrected) of the correlation of theta-activity during the probe trial with memory performance (average distance to target during probe trial); brain areas contributing to the significant correlation encompassed the temporal lobes, particularly in the right hemisphere and including the hippocampus and parahippocampal area, inferior occipital areas and/or cerebellum, as well as the right hemispheric striatum.

Figure 3C and D show the same for the correlation of average distance to target during the Probe trial and EEG activity in the immediately preceding trial 1, i.e. during encoding. Hence, here the EEG activity is not from the very same period during which the participant was moving across the maze, but it reflects neuronal activity of encoding, upon which the behavioural success of the probe trial depends. Note the remarkable similarity of the spectral and spatial pattern of the significant cluster (p < 0.05) to **Figure 3A and B**, despite the EEG data being taken from a different period than the behavioural measure.

To verify the specificity of these neuronal activation patterns to encoding and retrieval of one-trial place memory, we conducted three further control analyses:

1. We used the same average distance to target during the probe trial and correlated this to brain activity during corresponding trials 4; this failed to reveal any significant clusters (all p > 0.3, positive or negative).
2. We then correlated the average distance to target from all trials (initially including probe trials) to their corresponding EEG activity. This revealed a significant cluster in occipital and temporal areas with a similar spectral profile (p < 0.05, not shown) and spatially not too dissimilar to Figures 3A-D. No positive clusters were found.
3. We repeated this analysis, but this time removing all probe trials, and thus correlating average distance from target to simultaneous EEG activity in trials 1, 3 and 4 of the second locations of each block (i.e. loc. 4, 6 and 8) and trials 1, 2, 3 and 4 from the first location of all blocks (3, 5 and 7) which never contained a Probe trial, see Fig. 1B). This revealed a cluster that is, for comparison to **Figures 3A-D**, depicted in **Figures 3E** and **F**, showing a clear maximum in occipital cortex and a spectral peak in the alpha-band – but that did not reach omnibus significance (p = 0.14). From this dissociation we concluded that the highly similar outcomes shown in **Figures 3 A-D** were specifically reflecting processes related to encoding and retrieving/expressing memory representations as behaviourally measured in terms of search preference during the probe trial.

### Positive correlations of alpha-oscillatory activity with distance to target – potential correlates of motor behaviour

The cluster-permutation algorithm separates clusters showing positive and negative effects. In addition to the negative correlations of EEG activity with average distance to target, i.e. patterns reflecting increased activity with better memory performance, our analysis also revealed several positive correlations of oscillatory activity during probe trials (retrieval/expression) and during trial 1 (encoding) with average distance to target during probe trials (i.e., more oscillatory activity predicted lower search preference), which were specific with respect to frequency and brain region (**Figure 4**). **Figures 4A-B** show the positive correlation of alpha-activity (12 Hz) in motor cortex (left central sulcus over the hand-area) during the probe trial to average distance from target during the same probe trial, although this did not reach omnibus significance (p > 0.2). Moreover, as shown in **Figures 4C-D**, there was also a positive correlation for average distance to target with EEG activity during the (immediately preceding) trial 1, i.e. encoding. This positive correlation, with a similar alpha-/beta (12-14 Hz) peak and a location covering large parts of parietal cortex contralateral to the navigation hand and, thus, sensorimotor areas related to visually guided hand movements (Pause et al., 1989; Buneo and Andersen, 2006), was of omnibus significance (p < 0.05). For an interpretation of these correlations, see further below.

**Figure 4:**
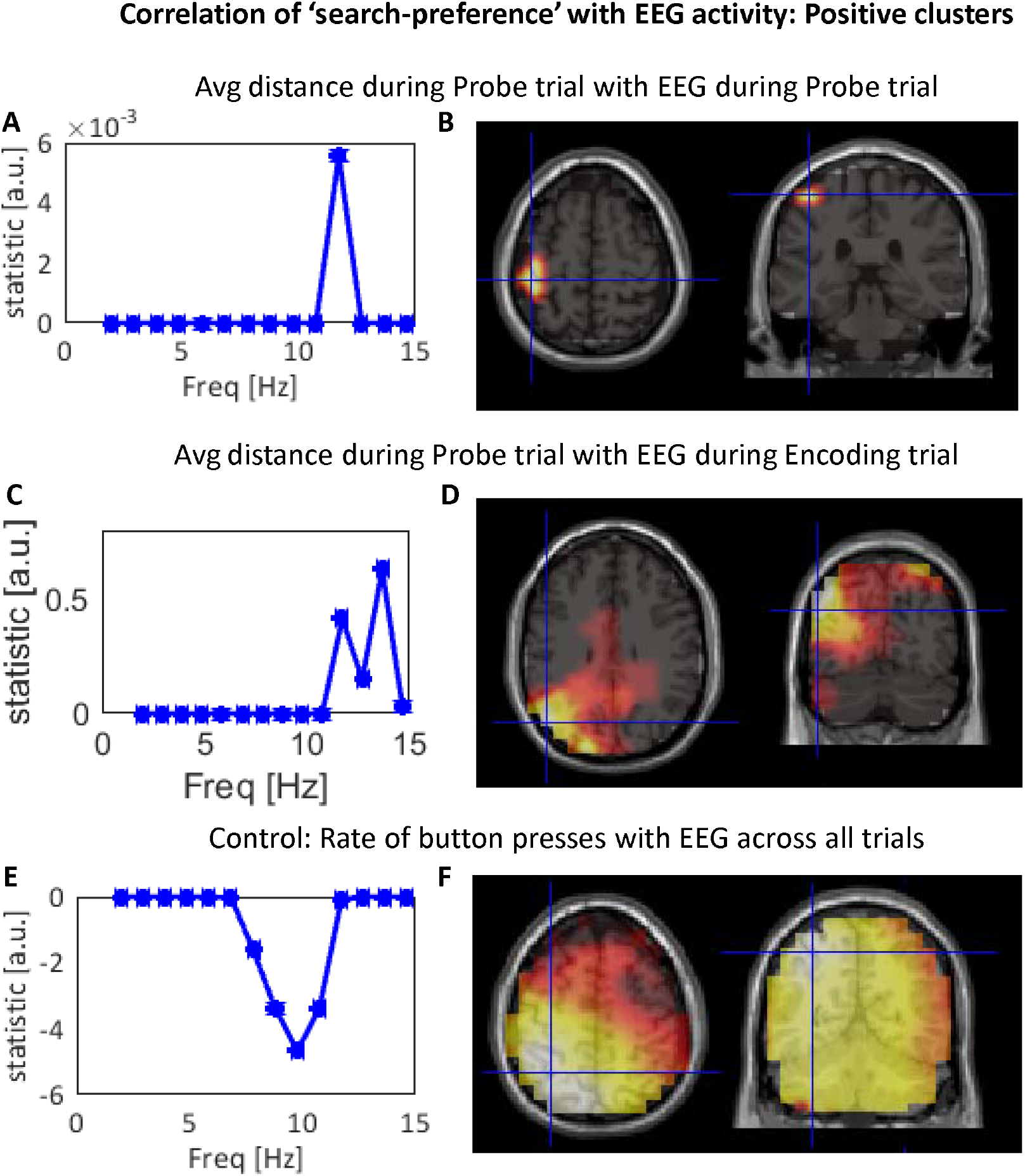
Positive correlations between oscillatory EEG activity and distance to target. **A-D:** Positive correlations of EEG activity during probe trials (retrieval/expression) and trial 1 (encoding) with average distance to target during probe trials. Crosshairs in brain topographies show the location of cross-section planes (chosen to provide good overview). **A)** shows the spectrum of this effect for the probe trial, whilst **B)** shows its topograpy with a clear maximum over left primary sensorimotor cortex. This cluster, however, failed to reach omnibus significance. **C)** shows the spectral distribution of the highly significant (p <0.01) cluster for the encoding trial 1, whilst **D)** shows its brain topography over left parietal areas. Since these effects, located over motor and sensorimotor regions, likely reflect motor-related patterns, a control analysis was conducted: The rate of button presses (see main text) was correlated with EEG activity across all trials. **E)** shows the spectrum of this effect. Note that a higher rate of button presses (i.e. less stationary movement patterns) correlates with reduced alpha-activity (whereas in A-D the average distance from target, a value likely to be inversely related to button-press-rate was used. **F)** shows the topography of this highly significant effect (p<0.001), which is similar though more widespread than the effect in B) and D).

The location of these positive clusters in motor cortex and parietal cortex, contralateral to the hand that participants were instructed to use for button presses to move through the maze, suggests that the correlations are movement related since alpha-activity in the motor cortex (frequently also referred to as mu-activity) is strongly associated with movement planning and execution (Pfurtscheller and Lopes da Silva, 1999). More specifically, they may reflect a) direct movement-related activations of the motor cortex contralateral to the hand that participants were instructed to use for moving and b) procedural or motor-learning that may facilitate retrieving the target location. We tentatively (and speculatively) interpret these two results therefore in the following way:

1. For the non-significant (when considering multiple comparison correction) correlation during the probe trial itself (where EEG data reflect the same period as the behaviour, Figure 4A-B), this likely reflects a motor confound such that participants who remain closer to the target show less stationary movement patterns (more frequent turns and stop-and-go manoeuvres), requiring more frequent button presses and thus more alpha-suppression; thus, according to this explanation, enhanced distance to the target would coincide with fewer button presses and thereby less motor-related alpha-suppression, i.e. higher alpha activity.
2. For the omnibus significant positive correlation of alpha-activity during the encoding trial 1, (da Silva et al., 2014) with average distance from target during subsequent probe trials (Figure 4C-D), i.e. impaired memory performance, we suspect that this may reflect that reduced movement (button presses), or exploration, during encoding results in weaker memory. This suggestion is consistent with the finding that the placement of rats onto the target location, instead of active swim trials, results in weaker memory in the watermaze DMP task (da Silva et al., 2014) and that volitional movements in human participants have been found to facilitate hippocampus-dependent memory in human participants (Voss et al., 2011).

In support of the implied relationships between movement during trial 1 or the probe, respectively, and average distance to target during probe trials, we found that the latter:

- correlated negatively with the rate of button presses during probe trials at r = −0.3 (p < 0.05); this means the closer participants remain to the target, the more frequently they press buttons on the same trial, probably because persistent searching close to the target location requires a high rate of button presses for frequent turns and stop and go (whereas continuous movement can be achieved by a low rate of button presses).
- correlated negatively with the rate of button presses during the preceding trial 1 at r=-0.2, albeit non-significantly (p > 0.05).
- correlated positively with the average speed of movement at r = 0.11, although this correlation was far from significance (p > 0.35).

To summarise this: 1) parameters of motor behaviour during the probe trial correlate significantly with the memory-related average distance to target and are likely causing the effect in sensorimotor cortex in **Figure 4A-B**; 2) more speculatively at this point, procedural or motor-learning and possibly motor parameters of exploratory behaviour (da Silva et al., 2014) during encoding in trial 1 may represent the correlations shown in **Figure 4C-D**.

This interpretation suggests that button presses across trials should negatively correlate with alpha activity. Indeed, an analysis using the rate of button presses (and thus the variable that correlated to average distance to target during probe trial) in all trials as a predictor variable for EEG activity in all corresponding trials revealed a highly significant (p < 0.001) negative correlation with alpha-activity, involving large parts (probably due to the effect size) of left parietal and motor cortex and of occipital cortex (**Figure 4E-F**). The direction of this effect is consistent with the well-established finding that alpha-activity is associated with activation of neural tissue (Pfurtscheller and Lopes da Silva, 1999), i.e. enhanced (sensori-) motor activity due to frequent button presses causing movement transients and therefore also non-stationary visual input. Importantly, no positive cluster was observed when button presses were used as predictor variable. Therefore, while movement-related factors are likely to account at least partially for the positive clusters in **Figure 4A-D**, there is no evidence that the memory network correlations presented in **Figure 3** are a result of movement confounds.

## Discussion

Combining the virtual DMP task with continuous EEG measures and using a correlational analysis unconstrained by a priori hypotheses on frequency bands or regions of interest, we have shown that better one-trial place memory performance (as reflected by average distance to target during probe trials) is predicted by theta oscillations within a largely right-hemispheric network consisting of MTL (including parahippocampal area), striatum and inferior occipital cortex or cerebellum, both during encoding and during retrieval/expression. Additionally, it emerged that enhanced memory performance was also correlated with reduced alpha activity in sensori-motor cortical areas contralateral to the navigation hand during encoding (left parietal cortex) or probe trials (left motor cortex, although this correlation was not statistically significant), probably reflecting movement-related factors that may at least partially be associated with memory performance.

### Comparison to previous MEG studies demonstrating hippocampal theta correlates of spatial memory and navigation

Previous studies (Cornwell et al., 2008; Kaplan et al., 2012; Kaplan et al., 2014; Pu et al., 2017) have used MEG in combination with individual anatomical MRIs to demonstrate hippocampal theta contributions during encoding (Cornwell et al., 2008; Pu et al., 2017) and retrieval (Kaplan et al., 2012; Kaplan et al., 2014) of spatial memory and navigation in virtual mazes. In the present study, the theta correlates of place memory performance were more pronounced in the right hemisphere (although some left temporal lobe activation was also evident), in line with hippocampal theta correlates reported by Kaplan and colleagues (2012, 2014) and Pu et al. (2017), and a wide range of evidence indicating that human spatial navigation memory is mainly associated with the right hippocampus (Burgess et al., 2002; Miller et al., 2018).

We, similar to (Cornwell et al., 2008; Pu et al., 2017), used an open-field navigation task, reverse-translated from a rodent watermaze paradigm, whereas (Kaplan et al., 2012; Kaplan et al., 2014) have relied on more structured tasks and used shorter epochs corresponding to specific encoding and retrieval periods to target temporal windows with an enhanced chance to detect hippocampal activation. Importantly, we used an analysis unconstrained by prior hypotheses on regions of interest or frequency bands to reveal theta originating from the MTL as a major neural mechanism underlying inter-individual differences in spatial learning. Interestingly, Cornwell et al found a correlate of theta activity in hippocampus to the reduction in path-length from trial 1 to trial 2. In the present study, we did not find an omnibus significant oscillatory correlate of reductions in path lengths measures from trial 1 to 2. However, note that: 1) in our data there was relatively small inter-individual variation in the difference from trial 1 to trial 2, as can be appreciated by the small error-bars in Figure 2; and 2) we may have used a more conservative statistical approach, also to deal with increased leakage of our spatial filters in EEG recordings based on a canonical forward model. Moreover, previous studies in humans and rats suggest that reductions in path lengths measures from trial 1 and 2 and search preference on the DMP task may rely on partially dissociable neuropsychological mechanisms (Buckley and Bast, 2018). First, there is only a moderate correlation between these measures in both human participants and rats (Buckley and Bast, 2018). Second, in rats, only the search preference measure shows a significant decline with increasing retention delay between trial 1 and 2, reflecting forgetting (da Silva et al., 2014). Third, and most importantly, studies in rats show that the search preference measure is more strongly hippocampus dependent, being more sensitive to disruption of hippocampal function by partial hippocampal lesions (Bast et al., 2009) and pharmacological manipulations (Pezze and Bast, 2012; McGarrity et al., 2017). Our finding that only the search preference measure showed theta correlates in the MTL is consistent with these previous findings.

### Reliability of our source-reconstruction

Analysing frequencies from 2-15 Hz, as a trade-off between statistical power/sensitivity and ensuring a broad range, we were able to confirm – whilst correcting for multiple comparisons in 28,120 data points in frequency and spatial domain – that it was predominantly theta-oscillations, in particular in the temporal lobe, that gave the best prediction of individual one-trial place memory performance on the virtual DMP task. Moreover, encoding and retrieval/expression correlates of memory performance were highly consistent, an issue we shall come back to further below. Although the anatomical locations of the theta correlates have to be interpreted with some caution on the basis of methodological limitations, this consistency between encoding and retrieval inspires a certain amount of confidence in their accuracy and spatial precision. Regarding the methodological constraints, two aspects should be mentioned:

1. These results were obtained on the basis of a canonical (standard) forward model, based on the MNI brain and, therefore, based on averaged electrode-positions for all participants (since individual positions on a standard brain and head surface were deemed meaningless); this imposes significant limitations on the spatial specificity, but perhaps also sensitivity of the beamformer solution (Troebinger et al., 2014). To address this issue, we applied a relatively high (noise-) regularization of the covariance matrix. The comparably high spatial specificity of the functional images shown are also due to these showing a statistic that is thresholded for datapoints that survive a non-parametric significance threshold of p < 0.01 and form part of a cluster that crosses the omnibus significance threshold (for details please refer to (Maris and Oostenveld, 2007)).
2. It would not be formally correct to ascertain on the basis of these results that a particular data point, which forms part of the omnibus significant cluster, drives this effect. For instance, the fact that the right striatum forms part of the cluster does not imply that this region is involved in spatial navigation memory at a confidence level given by the significance threshold corrected for multiple comparisons. This confidence level only applies to the entire cluster, not to each individual data point contributing to the shape of the cluster. However, the fact that a) we chose a fairly conservative cluster significance criterion of p < 0.01 and b) we see strikingly similar activation patterns in two largely independent (for the part of the EEG data) comparisons, for encoding and retrieval, does instil confidence in these findings.

Finally, participants’ movement and the dynamic visual input they received during task performance (as participants were moving relatively freely through the virtual environment) may have influenced our findings. However, it is highly unlikely that the theta correlates of one-trial place memory performance are explained by movement or dynamic visual input factors alone. None of the control analyses on movement parameters, nor a control analysis of our behavioural predictor variable of interest (average distance to target during probe trials) on unrelated trials yielded remotely similar theta correlates (see also next section on the alpha-correlates in sensorimotor regions of the brain). Hence, we conclude that movement or dynamic visual input effects are not the main factors to the effects we see in the putative network of temporal lobe, striatum and inferior occipital and/or cerebellar activation. Encoding of spatial memories in the real world also requires movements and dynamic visual input, such that avoiding this would be undesirable.

### The brain network supporting one-trial place memory performance

Beyond the hypothesis-conform activation of the temporal lobe, including the hippocampal and parahippocampal region, we see further distinct activations in the right striatum and inferior occipital cortex and/or cerebellum – all at theta frequencies with a peak at 7 Hz. Activity in these regions, at this frequency, correlated negatively with distance from the target – suggesting that higher theta activity in these regions during both encoding and retrieval led to better one-trial place memory performance across individuals. Occipital involvement is consistent with the dependence of the task on visual input, and with previous studies implicating occipital theta activity in performance of virtual spatial navigation tasks (Ekstrom, 2015) and in working memory tasks in both humans and animals (Lee et al., 2005; Fuentemilla et al., 2014).

With respect to cerebellum and striatum, it is evident that the quality of the anatomical forward model used in the present study requires some caution and standard forward models developed for the cortical sheet are less suitable to detect such deep sources with cell geometries that may differ from the cortical sheet (Meyer et al., 2017). Nevertheless, our finding of a putative involvement of cerebellar theta activity adds to an emerging literature that the cerebellum may provide input on self-motion and trajectory learning into the hippocampus, to facilitate the establishment of a cognitive map (Lefort et al., 2015), and is consistent with the suggestion that interactions between MTL and cerebellum may be coordinated by theta activity (Watson et al., 2019). More directly, in one of the first, if not the first publication on electrophysiological hippocampal activation measured in humans in the context of a watermaze-like virtual maze, Cornwell et al. (Cornwell et al., 2008) showed a highly similar cerebellar theta activation in their MEG recordings.

The striatal correlates observed in the present study, in two separate conditions, are also consistent with some previous finding. Studies of human spatial navigation in a virtual maze, have implicated the striatum and hippocampus (Doeller et al., 2008), and dopaminergic decline in the striatal-hippocampal circuit during ageing and Parkinson’s disease has been associated with deficiencies in spatial navigation and memory (Thurm et al., 2016); these studies suggested that striatal mechanisms mainly serve cue-response strategies supporting spatial navigation, which are distinct from and complement hippocampal mechanisms of allocentric place representations. In addition, the human striatum has been implicated in performance on laboratory tests of hippocampusdependent declarative memory (Scimeca and Badre, 2012). Moreover, rodent studies that combined behavioural testing with multi-site electrophysiological recordings (Martin and Ono, 2000; DeCoteau et al., 2007; van der Meer and Redish, 2011) or disconnection approaches (Floresco et al., 1997; Devan and White, 1999) support that hippocampal interactions with the striatum, especially mediodorsal and ventral striatum (the main recipients of hippocampal projections; (Voorn et al., 2004; Humphries and Prescott, 2010)), are important for rapid place learning performance (reviewed in (Bast, 2011)). Studies in freely behaving rodents have also shown theta oscillations in the striatum and suggested that interactions between hippocampus and striatum, again mainly mediodorsal and ventral parts, are coordinated by theta oscillations in the context of spatial memory and navigation tasks (Berke et al., 2004; DeCoteau et al., 2007; Tort et al., 2008; van der Meer and Redish, 2011). Finally, in a recent study in rats, we found that functional inhibition targeting the ventral striatum disrupted one-trial place memory performance on the watermaze DMP task (Seaton Aetal., 2019).

The idea that theta-oscillations support memory and particularly enhance the precision of spatial memory representations has a long history. (Jensen and Lisman, 2000) for instance, found that when analysing the spike-time-series of place-field recordings, considering the position of the spikes in the phase of ongoing theta-oscillations substantially boosted the decoding accuracy of the rat’s position in the maze. (Lee et al., 2005) found, in a visual working memory task, that during the encoding period the delay activity was not characterised by an increase of spiking activity, but, instead, the spikes of those neurons carrying the stimulus representation strongly synchronises to the ongoing theta rhythm; this effect appears to be part of a network effect where phase-synchronization of spikes in visual cortex to a prefrontal cortex driven theta rhythm appears to be a major determinant on the success of working memory material retention (Liebe et al., 2012). Regarding the computational role of theta-oscillations, one prominent idea is that theta-oscillations facilitate and orchestrate spike-time-dependent plasticity (Lengyel et al., 2005; Sejnowski and Paulsen, 2006); theta-oscillations in different brain regions may ensure that the inputs from different neuronal populations may be timed such that effective spike-time-dependent memory traces can be formed and retrieved. Neurophysiological studies and computational modelling suggest how theta-oscillations in the hippocampus might play a mechanistic role during both encoding and retrieval (Hasselmo and Stern, 2014).

Overall, our study suggests that one-trial place memory performance may be supported by a brain network, coordinated by theta activity and including the MTL, occipital cortex, cerebellum and striatum. The particular nature of the interactions between these regions in the theta range would perhaps be more suitably studied with further studies utilising individual and specialised forward models in the context of MEG recordings (due to enhanced MEG specificity to study neuronal interactions between different brain areas).

### Sensorimotor correlates of DMP task performance

Beyond the theta-band effects, we also observed that our measure of search preference during probes correlated with some activity in higher frequency bands, notably the sensorimotor alpha-band (Pfurtscheller et al., 1996), in sensorimotor areas contralateral to the hand that participants were instructed to use for navigation by button presses. This correlation was (omnibus) significant only in correspondence to the encoding trial and revealed a relatively broad cluster of parietal regions. For EEG activity during the probe trial (retrieval), this did not reach omnibus significance, but was localised very clearly to the hand-area of the central sulcus (notably the critical p-value for individual data-points was p < 0.01). Control analyses considered in the Results section suggested that these correlates reflect movement-related factors associated with task performance. The nonsignificant correlation of alpha-activity in the hand area during the probe trials with search preference during the probe trials likely reflected that participants with more precise spatial memory utilised more button presses to perform frequent turns and stop and go. The significant correlation of alpha-activity in parietal sensorimotor regions, which have been implicated in visually guided hand movements (Pause et al., 1989; Buneo and Andersen, 2006), during trial 1 with search preference during the probe trials may reflect that increased active exploration, or enhanced attention to one’s movements (which would be expected to result in reduced alpha-power (Jensen and Mazaheri, 2011; Bauer et al., 2012a), during trial 1 boosts one-trial place memory. This suggestion is consistent with the finding in rats that placement of rats into the target location, instead of active swim trials to the target location, results in weaker memory in the watermaze DMP task (da Silva et al., 2014) and that in human participants volitional movements have been found to facilitate hippocampus-dependent memory (Voss et al., 2011).

### Conclusion and future directions

We have shown that, even when using a basic canonical forward model, surface EEG recordings in combination with the virtual DMP task can reveal the approximate neuroanatomy of a spatial memory system in the human brain on the basis of inter-individual variation in memory performance. This approach holds great potential for future studies on the potential oscillatory correlates of inter-individual differences, including in relation to age-related changes in spatial memory (Driscoll et al., 2005; Hort et al., 2007; Lester et al., 2017; Coughlan et al., 2018) with a task that is suitable for different age groups (our own unpublished work with elderly participants and previous work using a related task in children (Buckley et al., 2015)). Of particular interest will also be the combination with neuropharmacological techniques to study the neurochemistry of such brain oscillations (Newman et al., 2012), as well as their importance for inter-individual differences in healthy (Richter et al., 2014) and clinical populations (Meng et al., 2018). The comparably low costs and the availability of EEG recordings may make this also suitable as a diagnostic tool for clinical practice. Regarding the question of the more mechanistic role of theta-oscillations for interactions between different parts of the (putative) network that was revealed by our source analysis, these may be more suitably be addressed by MEG studies with sophisticated forward models (Meyer et al., 2017).

## Acknowledgements

We would like to thank Maazah Muhammad Ali, Zaariyah Hasan Bashir, Jessica Burrington, Irina Surducan, Abigail Webb and Desislava Abu Rashed for assistance with data collection. Markus Bauer was supported by a Nottingham Research Fellowship.

